# Deep learning-based event classification of mass photometry data for optimal mass measurement at the single-molecule level

**DOI:** 10.1101/2025.06.21.660868

**Authors:** Kishwar Iqbal, Jan Christoph Thiele, Dominik Saman, Jack S. Peters, Stephen Thorpe, Samuel Tusk, Jack Bardzil, Justin L.P. Benesch, Philipp Kukura

**Affiliations:** The Kavli Institute for Nanoscience Discovery, University of Oxford, Dorothy Crowfoot Hodgkin Building, South Parks Road, Oxford OX1 3QU, UK; Physical and Theoretical Chemistry Laboratory, Department of Chemistry, University of Oxford, South Parks Road, Oxford OX1 3QZ, UK

## Abstract

Mass photometry (MP) is a powerful technique for studying biomolecular structure, interactions, and dynamics in solution. It detects and quantifies small reflectivity changes at a glass-water interface during protein (un)binding, with signals typically averaged over 100 milliseconds. However, particle motion at the point of single-molecule measurement can compromise key metrics such as mass resolution, sensitivity, and concentration. We present a three-dimensional convolutional residual network trained via supervised learning to classify landing events based on their spatiotemporal dynamics. By analysing 3D event thumbnails, our method isolates optimal single-molecule measurements, eliminating cumulative histogram artifacts and improving resolving power by up to a factor of 2. Validated across diverse experimental datasets—including resolved and partially resolved samples, and varying masses, concentrations, and integration times—our approach delivers robust performance under (sub)optimal conditions. Our approach provides measurement-level data-driven feedback, facilitating high quality MP measurements in challenging scenarios.

## Introduction

Mass photometry (MP)^1^ enables label-free studies of biomolecules with single-molecule sensitivity. By accurately measuring the mass of individual molecules and their complexes in solution, MP enables precise quantification of oligomerisation, protein–protein, and protein–DNA interactions, as well as complex assembly^2–10^. Typically, a sample is added to a microscope coverslip (**Fig. 1A**), where non-specific binding to the glass occurs. Because the refractive index of biomolecules (e.g. proteins, ∼1.46) differs from that of water (1.33), each binding event produces a local change in reflectivity. This change scales with molecular polarizability, which depends on the refractive index difference between the biomolecule and its surroundings, and is proportional to its volume. The similarity in optical properties and density among biomolecules results in a direct proportionality between polarisability and mass, enabling precise mass measurements at the single molecule level.

**Fig 1.**
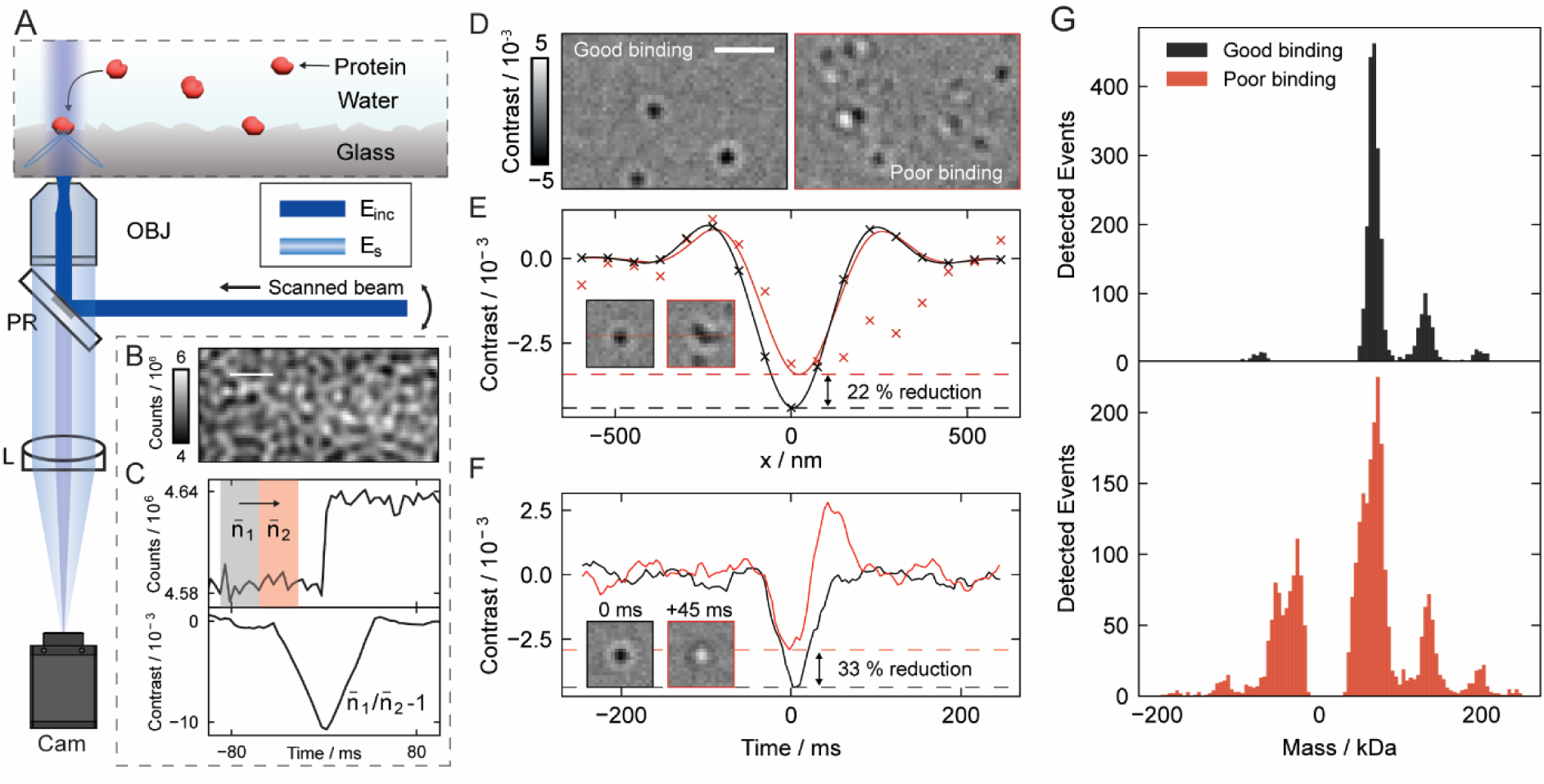
Mass photometry and challenges for accurate mass quantification. (A) Simplified schematic of a mass photometer and protein landing assay. (B) The raw camera image is dominated by scattering from the cover glass roughness. (C) Ratiometric image processing, achieved by dividing sequential frame substacks (𝑛_1_and 𝑛_2_), removes the background, revealing the weak signal from single-molecule landing events. (D) Images depicting (sub)optimal binding of proteins to the glass surface. (E) Interference from overlapping molecules can lead to signal quantification inaccuracy (22% reduction shown). (F) Rapid unbinding within the ratiometric processing window can also result in a signal reduction (33%). (G) – The cumulative impact of many inaccurately quantified events can introduce artifacts in the mass histogram, as shown in measurements of fresh BSA stock (black) and old stock (red). Abbreviations: PR – Partial Reflector, OBJ – Objective, L – Lens, Cam – Camera, E_inc_ – Incident field, E_s_ – Scattered field. Scale bar: 1 µm.

When experimental and shot noise are sufficiently reduced, these reflectivity changes can be visualised in real time, even against a substantial optical background (**Fig. 1B**). In cases of irreversible binding, the signal appears as a step function (**Fig. 1C**) which is traditionally processed through a ratiometric analysis (**Fig. 1C, 1D**)^11,12^. To enhance precision, the recorded point spread function (PSF) is fitted to an analytical model (**Fig. 1E, line**), yielding the contrast that, when compared to a calibration, yields the molecular mass. The precision of these single-molecule mass measurements depends on background noise, the shot noise contribution to which can be reduced by temporal averaging of frames recorded before and after binding (𝑛̅_1_, 𝑛̅_2_). However, this averaging is only successful and representative if binding is irreversible and unperturbed during the integration time - referred to as *optimal binding*. For example, the binding of a second, neighbouring molecule (<1 µm, <100 ms) distorts the PSF, resulting in an inaccurate mass estimate (**Fig. 1E, red**). Similarly, transient unbinding can cause interference between binding and unbinding signals within the integration window, leading to contrast underestimation and mass error (**Fig. 1F, red**). Other perturbations such as lateral movement due to imperfect binding—whether simple translation ("rolling") or erratic oscillations ("wobbling")—further disrupt the PSF and thus quantification accuracy. Recording and quantifying all of these events generates a mass histogram (**Fig. 1G**). However, a substantial presence of these suboptimal events can distort the histogram features, causing loss of resolution through mass broadening, artificial low-mass peaks, artifacts at negative mass from unbinding, and increased baseline noise from poorly quantified low-abundance events (**Fig. 1G, red**).

To quantify resolution in mass photometry, we adopt the term *resolving power*, defined analogously to mass spectrometry as 𝑚/𝛥𝑚^13–15^, where *m* is the mass and *Δm* the specified peak width. For isolated peaks, we take the full width at half maximum (FWHM) to determine 𝛥𝑚. In cases of substantial peak overlap, we use the valley definition, where 𝛥𝑚 corresponds to the mass difference between two peaks of similar height such that the valley between them drops to a specified percentage (e.g., 10%) of the smaller peak. Importantly, this definition captures a mass-dependent, quantitative measure of MP performance, allowing consistent comparisons across samples and experimental conditions. Advancing the resolving power is as crucial for mass measurements as improving resolution has been for structural biology^16^. It determines whether molecular species in increasingly complex mixtures can be distinguished or remain unresolved, thus transforming the applicability of the technology.

MP has gained rapid adoption due to its ease of use and ability to quantify mixtures. Poor binding remains a challenge, because it impacts key performance metrics—resolving power, sensitivity, and concentration range—complicating data interpretation. Surface functionalisation can mitigate this issue but often lengthens and complicates experimental protocols^17,18^. If a measurement contains a sufficient number of optimal binding events, they can in principle be extracted through analytical techniques such as residual filtering. However, these methods are used on a case-by-case basis and often require expert knowledge, are time-consuming, and can become subjective.

The introduction of deep learning has revolutionised the field of data-driven analysis^19–28^. It has been widely applied across various biophysical techniques, including fluorescence-based microscopy^29–32^, mass spectrometry^33,34^, and cryo-EM^35–37^, and more recently, in interferometric scattering microscopy ^38,39^. These techniques have yet to be explored in the context of mass photometry performance — crucial to enable quantification of progressively more challenging biomolecular samples and mixtures.

In this work, we implement a three-dimensional convolutional residual network trained via supervised learning on a dataset of simulated surface binding events over an experimentally acquired background. The model classifies each single-molecule binding event based on its local spatiotemporal features, represented as a 3D-thumbnail input, enabling the identification and retention of high-quality measurements. We evaluate our model across diverse and representative test conditions, including optimal and suboptimal binding measurements, varying mass and concentration levels, and different integration times—validated on over 100,000 experimental and 3 million simulated events. We demonstrate our model’s ability to refine experimental data by correcting the mass histogram and eliminating artifacts caused by inaccurate quantification. Notably, we show improvements in resolving power by up to a factor of 2. The most challenging test case was heat shock protein 27 (HSP27, bird), a poorly binding highly polydisperse protein. Our model substantially increased the resolution enabling clear separation of previously unresolved species, with validation through native mass spectrometry. Additionally, our approach introduces a new layer of quantitative feedback by revealing the distribution of optimal and suboptimal events within a given measurement. This feedback enables more consistent, high-quality MP measurements at the technique’s performance limits, helping to unlock more complex systems that remain challenging for MP.

## Results

### Quantifying the effect of suboptimal binding

We broadly classify MP landing events into five categories: **‘**binders**’** (optimal) and four suboptimal types—**‘**unbinders**’, ‘**neighbours**’,** ‘rollers’, and ‘wobblers’, as illustrated in **Fig. 2A**. These classifications are guided by domain knowledge derived from extensive analysis of experimental data, capturing recurring behaviours that commonly effect mass accuracy. To assess the impact of suboptimal binding on single molecule mass measurement performance, we simulated landing events for a 180 kDa protein under varying conditions (**Fig. 2B**), adjusting parameters such as binding duration, unbinding probability, and lateral velocity to capture different landing dynamics (**Table S1**, **Fig. S1**). Our analysis reveals that 85% of optimally binding events achieve a mass accuracy deviation of less than 5%, compared to only about 30% of suboptimal events reaching this level of precision. This inaccuracy distorts the mass histogram, leading to broadening of peaks in the mass distribution, with a direct impact on resolving power, and an increased noise floor (**Fig. S2**). Analysing fit residuals provides a means to distinguish spatially dissimilar events, such as neighbouring or rolling events. However, it fails to differentiate transient behaviours, such as rapid unbinding or wobbling, which require temporal information. Residuals also lack interpretability, leaving users unaware of why certain events yield poor quantification.

**Fig. 2.**
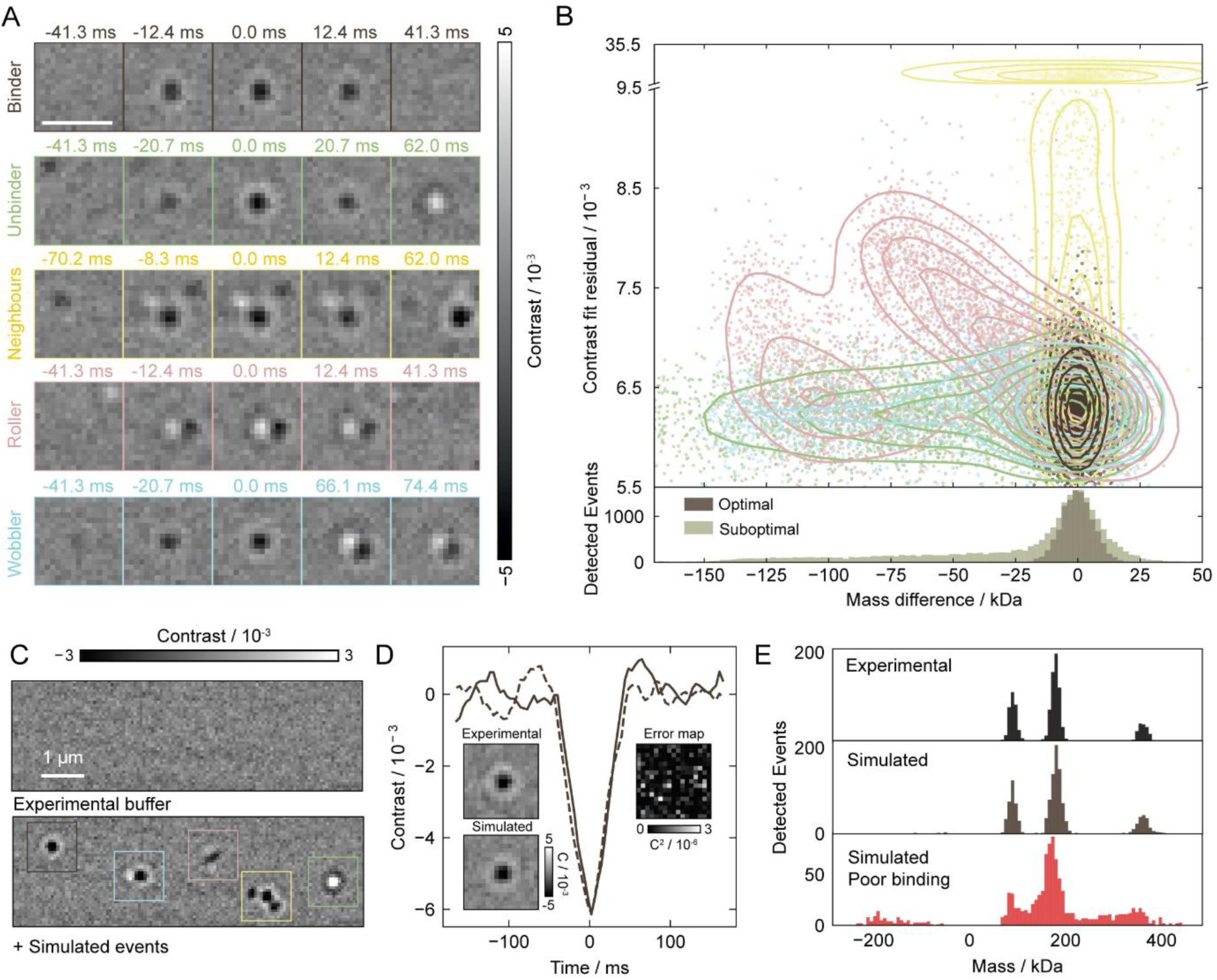
Analysis and simulation of the impact of suboptimal binding. (A) - Five common landing event types in mass photometry. (B) - Relationship between mass quantification accuracy and fit residual for 25,000 simulated 180 kDa molecules. Optimal quantification is achieved for stably binding molecules, while all other event types introduce increased mass inaccuracy, primarily leading to underestimation. Counts are displayed below, with optimal event counts normalised for comparison. (C) - Simulation framework using synthetic particle signals with diverse landing dynamics, overlaid on an experimental background. (D) - Comparative analysis between an experimental (solid) and simulated (dashed) event, demonstrating high agreement, as seen in the error map. (E) - Comparison between experimental and simulated mass histograms.

This analysis was enabled by our ability to accurately simulate MP data. We achieve this by modelling each binding event using experimentally determined PSFs superimposed on experimental background noise extracted from MP movies of buffer blanks without analyte (**Fig. 2C**). To validate this approach, we compared experimental (solid) and simulated (dashed) events, demonstrating close agreement (**Fig. 2D**). As a result, we can generate complete MP movies that produce mass histograms virtually indistinguishable from experimental data, both for optimal and suboptimal binding (**Fig. 2E**). Notably, this simulation framework also serves as the foundation for our supervised learning training dataset (see Methods for details).

### Deep learning-based framework

We generated a dataset of 25,000 simulated landing events, evenly distributed across event classes with varying parameters to capture a broad range of spatiotemporal characteristics (see Methods for details). As a first proof of principle of this framework, we simulated events spanning the 30–800 kDa mass range, with a skewed emphasis on the 30–100 kDa mass range, where the lower signal-to-noise (SNR) ratio presents greater challenges for accurate mass measurement. The dataset was split 80:20 for training and validation to optimise model learning while ensuring reliable generalisation (**Fig. 3A,B**). We implemented a 50-layer 3D convolutional residual network based on the ResNet architecture (**Fig. 3C**)^21,40^ to classify different types of landing events. Residual connections were integrated at the start of each residual layer, enabling deep feature extraction crucial for enabling separation between subtly wobbling events from optimally binding ones, particularly in low-SNR measurements (**Fig. 3D,E**). The model receives the local spatiotemporal information of each landing event through the input of a (40 frames by 17 x 17 pixels) thumbnail, centred around the detected landing event. It then assigns class scores to determine the event type.

**Fig 3.**
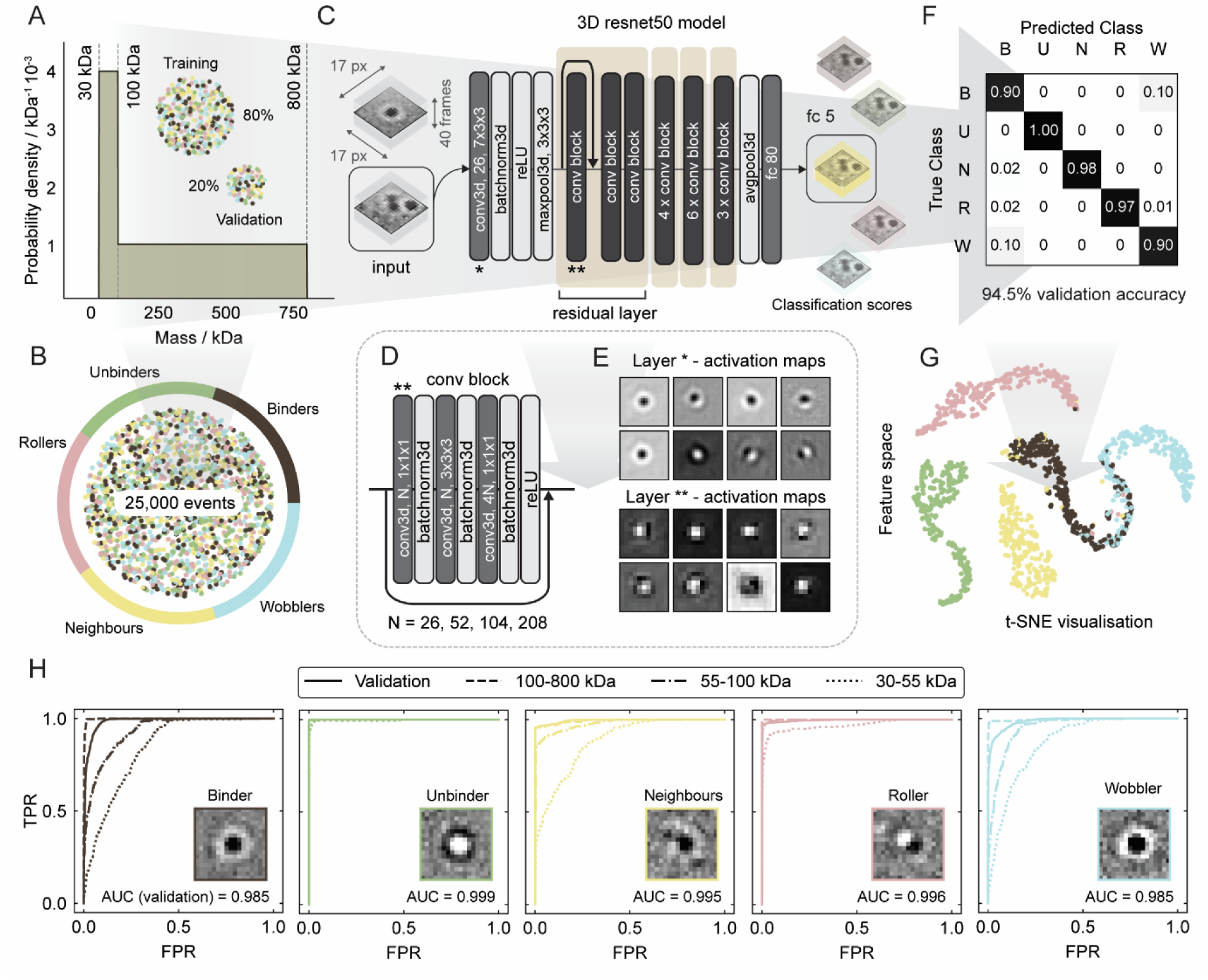
Deep learning-based event classification: An overview. (A) - Mass distribution of the training and validation dataset (80:20 split), with emphasis on the low signal-to-noise ratio range (30–100 kDa). (A) (B) The dataset was generated with a balanced distribution of 25,000 simulated events across event types. (C) - A 50-layer 3D convolutional residual network classifies single-molecule landing events based on their local spatiotemporal dynamics, using a (17 px by 17 px by 40 frames) event thumbnail as input. (D) The network consists of residual connections after the first convolutional block in each residual layer. (E) A temporal slice from the 3D activation maps illustrates the model’s response in the first two convolutional layers. (F) - Confusion matrix from the validation dataset, highlighting overall classification accuracy. (G) t-distributed Stochastic Neighbour Embedding (t-SNE) visualisation of the event feature space, illustrating the separation between the different predicted classes. (H) - ROC curves illustrating diagnostic performance by class on the validation dataset, along with additional simulated datasets (1,500 events each) segmented by mass range. The model shows better performance for masses above 100 kDa.

After training (see methods), our model achieved 94.5% validation accuracy. The confusion matrix revealed that binding and wobbling events were most frequently misclassified (**Fig. 3F**), highlighting the ambiguity in separating events with very small perturbations. This trend was further reflected in the t-SNE visualisation, which showed substantial overlap between these two event types (**Fig. 3G**). We further evaluated classification performance per class across the 30 to 55 kDa, 55 to 100 kDa and 100 to 800 kDa mass ranges using receiver operating characteristic (ROC) curve analysis (**Fig. 3H**). **Table S2** presents the area under the curve (AUC) for each test case, demonstrating strong overall performance. Notably, high classification accuracy is maintained across a broad mass range, however, this becomes more challenging near the detection limit (30–55 kDa) due to the low SNR.

### Event Classification for Mass Photometry

We evaluated our model on experimental data with a variety of protein mass distributions and binding affinities to glass that are representative of the most often faced challenges for MP measurements (**Fig. 4**). In a first application, we retained only optimal binding events to enhance MP performance. We set the binding score threshold based on the ROC curve analysis on the validation data, while balancing the need to retain sufficient counts per peak after filtering. We measured BSA as a reference protein, where both optimal and suboptimal binding are observed in independent measurements depending on sample quality (**Fig. 1**). The model accurately classified events from the mixed BSA dataset, as confirmed by inspection of the single-molecule measurements (**Fig. 4A**). The suboptimal events were primarily responsible for counts appearing between BSA peaks due to erroneous mass quantification. These artifacts are often mirrored on the negative mass side of the histogram, as unbinding events. By selectively removing these poor events, the model effectively eliminates artifacts introduced by suboptimal landings. Importantly, the ratio between different oligomeric states is essentially unaffected. In cases where low-mass species may be underrepresented due to SNR-related misclassifications in mixed samples, this bias can be estimated and corrected (**Fig. S3**).

**Fig 4.**
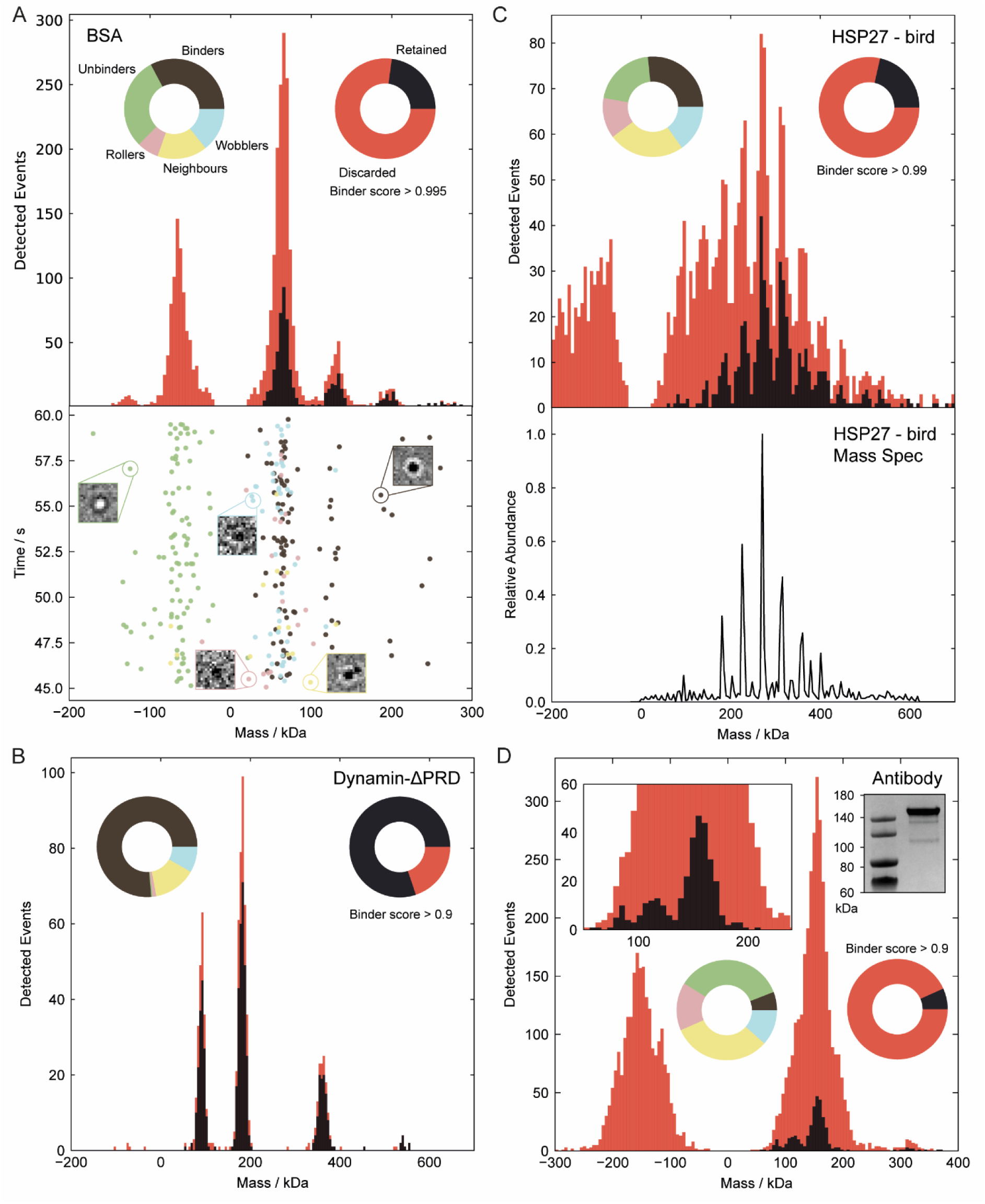
Identifying optimal single-molecule landing events for diverse test protein measurements. (A) Bovine serum albumin. Pie charts display the predicted event class distribution and selective retention based on the optimal binder score threshold. The mass histogram is shown before (red) and after (black) event filtering, with the distribution of events over time during the final 15 seconds of the landing assay shown below. (B) Dynamin-ΔPRD. (C) Heat shock protein 27 (HSP27). Mass distributions before and after filtering are shown (top), with validation provided by native mass spectrometry (bottom). (D) Anti-PSMC6 antibody, with validation provided by SDS-PAGE analysis (inset).

By contrast, dynamin-ΔPRD measurements represent a protein with high binding affinity to glass and exhibits predominantly optimal binding events. Here, the stochastic nature of landing events on glass and the event density led to neighbouring events being the highest proportion of discarded events. This resulted in a largely unchanged mass distribution before and after filtering (**Fig. 4B**). This represents the expected level of optimal performance, corresponding to a resolving power of approximately 10 for the 180 kDa dimer peak and ∼17 for the 360 kDa tetramer peak HSP27 represented the most challenging protein we tested, due to its broad mass distribution spanning much of the mass range over which our model was trained. It also presented many partially resolved peaks and exhibited poor surface affinity. Event classification and filtering revealed that the inclusion of suboptimal events (see **Supplementary Video 1**) not only elevated the baseline of the mass distribution but also artificially increased counts at the low-mass end. After their removal, a near-baseline resolved distribution was achieved and validated through native mass spectrometry, as shown in **Fig. 4C**. We quantified the resolution improvement using the valley-to-peak ratio of the partially resolved peaks, as reported in **Table S3**. These were also compared to standard and user optimised results from the Discover^MP^ software (Refeyn). Prior to filtering, the oligomeric peaks of HSP27 could not be resolved under the 30% valley criterion. Optimising the analysis parameters in Discover^MP^ allowed the 8–12-mer peaks to marginally meet this threshold. However, arbitrary adjustments to Discover^MP^ parameters can alter the mass distribution and lead to incorrect results, highlighting the need for expert tuning (**Fig. S4**). Following our filtering approach, all peaks were resolved using the 30% criterion, with the 6–10-mer peaks even satisfying the more stringent 10% condition—demonstrating a substantial improvement in resolution. Residual analysis shows our method removes both high-residual outliers and low-residual inter-cluster events that broaden peaks and raise the baseline for HSP27 (**Fig. S5**). Overlaying mass and residuals in t-SNE space highlights the largely mass-independent nature of our classification (**Fig. S6**).

For an anti-PSMC6 antibody sample, which exhibited poor surface affinity, the filtering process removed many events caused by significant unbinding from glass (**Fig. 4D**). These events resulted in a mirrored negative mass peak which also increased the event density, leading to many neighbouring events. Filtering substantially increased the resolving power of the 150 kDa peak from 3.9 ± 0.5 to 7.6 ± 0.3 across 8 independent measurements (29,447 total events), approaching the level expected for an optimal measurement (i.e., **Fig. 4B**). The enhanced resolution revealed low-abundance degradation products in the antibody sample, observed as peaks below 120 kDa which were validated by SDS-PAGE analysis (**Fig. 4D, inset**). We further tested our model for massference-p1 and apoferritin samples, where it effectively cleaned the mass histograms of artifacts from poorly quantified measurements (**Fig. S7**).

In addition to sub-optimal binding, excessive binding density can also contribute to inaccurate mass measurement. At low average event densities of 0.3 μm^-2^s^-1^, we found negligible changes to the resolving power of the tetramer peak of dynamin-ΔPRD at 360 kDa before and after filtering (**Fig. 5A-C**). Most events were retained due to the low probability of overlap and interference. As the event density increases with analyte concentration, the proportion of discarded events also increases due to an increase in overlapping events. As a result, the degree of improvement in mass resolution by our method increases with higher event densities. For example, at an event density of 34.6 μm^-2^s^-1^, the resolving power improved from 5.3 ± 0.3 to 10.0 ± 1 after filtering. Despite the observed improvement, the mass resolution at higher concentration remains limited compared to that achievable at lower concentrations due to a general increase in imaging background.

**Fig 5.**
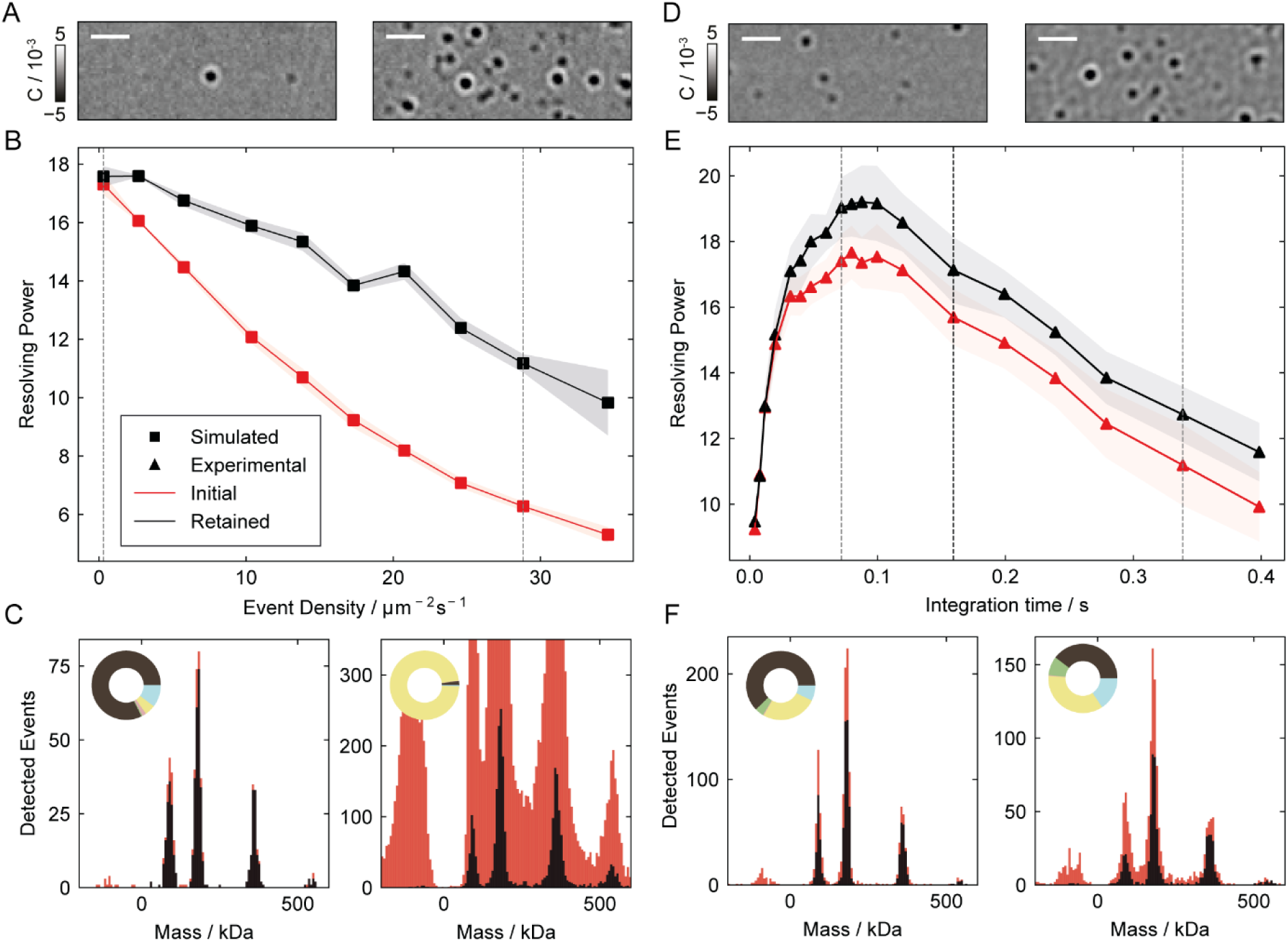
Optimal event selection improves resolution with increasing concentration and integration time. (Left) Effect of suboptimal event removal on simulated Dynamin-ΔPRD assays with increasing event density, with 5 repeats per concentration. (A) Frames for landing densities highlighted by the dashed lines in B. (B) Resolving power as a function of event density for the 360 kDa tetramer species, plotted before (red) and after (black) selective filtering. (C) Mass histograms corresponding to the highlighted densities. Binder score threshold was set to 0.8. (Right) Effect of event filtering on experimental Dynamin-ΔPRD assays analysed across increasing integration times, repeated for 5 measurements. (D) Frames analysed at 70 and 340 ms integration times, as highlighted by the dashed lines in E. (E) Resolving power as a function of integration time for the 360 kDa tetramer species, plotted before (red) and after (black) event filtering. The integration time was varied for the particle fitting process. Particle detection and classification were performed at two fixed integration times: 40 ms integration for data fitted below 160 ms, and 160 ms integration for data fitted above 160 ms. (F) Mass histograms corresponding to the highlighted integration times.

Increasing the integration time for a given analyte concentration also raises the probability of partially overlapping events. A longer temporal integration window also makes single-event measurements more susceptible to transient effects, such as rapid unbinding. We observe a clear improvement in resolving power of the tetramer peak of dynamin-ΔPRD as the integration time increases to 20 ms, attributed to the improving SNR of the measurements (**Fig. 5D-F**). In this range, the filtered and non-filtered data show minimal differences, owing to the short temporal window and low perceived event density. The increase in resolving power continues and reaches an optimal point between 80 - 120 ms. Within this range, we observe up to a 10% improvement, reaching 19 ± 1 following event filtering. This occurs due to the mitigation of the effect of overlapping/transient events. Beyond 120 ms, the mass resolution begins to decrease due to the increased density and incorporation of non-shot noise contributions. Although event filtering improves performance, it fails to achieve optimal levels. We partially attribute this to the model not being explicitly trained on experimental noise sources, such as those arising from the glass surface or fluctuations in the illumination.

We then evaluate the performance of our model near the quantitative detection limit at 40 ms integration with low-SNR measurements of protein A (42 kDa). For standard measurements, a low detection threshold required for low mass detection leads to detection of many false events, which manifests itself as a noise peak symmetric about zero mass in the mass histogram, illustrated on a buffer measurement (**Fig. 6A**) and for protein A (**Fig. 6B**, red). High detection thresholds, on the other hand, result in an asymmetric cut off on the low mass end of the detected peak (**Fig. 6B**, **inset** (red)). After filtering protein A events using our model (**Fig. 6B**, main panel and inset, **black**), we found a more symmetric peak centred around the expected mass of the protein (42 kDa), with false detections effectively removed. We applied a high classification threshold to retain only events with high certainty, however, this led to discarding approximately 70% of protein A events. Titration of protein A demonstrates that our approach remains sensitive enough to quantify changes in the protein A sample concentration without interference from spurious background detections (**Fig. 6C**). While not representing a substantial improvement in low-mass detection, this analysis offers a useful benchmark of model performance near the detection limit and informs its application to smaller proteins discussed later. We validated the discarded events from the noise peak by evaluating it at a 140 ms integration time. The higher SNR shows that these spurious detections did not correspond to real events (**Fig. 6D**). This becomes clear when comparing them to the retained events imaged at 140 ms integration (**Fig. 6E**).

**Fig 6.**
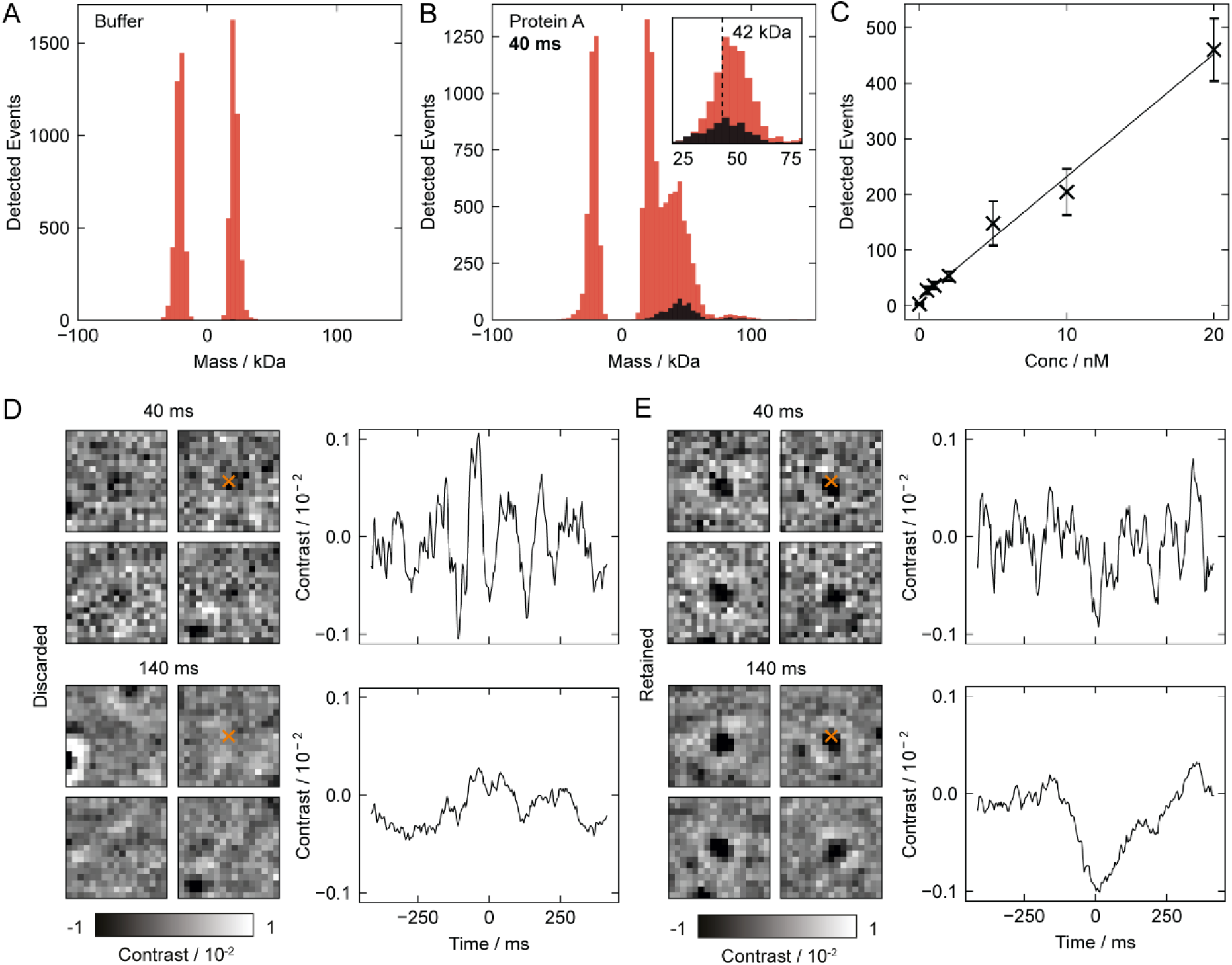
Detection of protein A (42 kDa) at 40 ms integration time. (A) Analysis of buffer blanks with low detection thresholds depicting false event detection. (B) Analysis of protein A under the same detection conditions. Selective event filtering was applied, with discarded events shown in red and retained events in black. The inset highlights the asymmetric peak (red) observed when applying high detection thresholds. (C) Titration of protein A counts. (D) Selected event thumbnails and time traces of discarded events, shown at 40 and 140 ms integration. (E) Selected event thumbnails and time traces of retained events, shown at 40 and 140 ms integration.

## Discussion

Precise quantification of single-molecule landing signals underpins the mass measurement and resolving capabilities of mass photometry, enabling its broad applicability. Suboptimal surface binding and interference from neighbouring molecules can distort signal quantification, limiting the sensitivity, resolving power, and concentration range of the technique. To address this, we adapted and optimised a 3D ResNet50 model to classify single-molecule landing events in MP based on their spatiotemporal features, enabling accurate separation and classification of optimal and suboptimal events. Applied across a diverse set of proteins varying in molecular mass, surface affinity, concentration, and integration time, our method removes artifacts and improves resolving power by up to a factor of two, recovering near-optimal performance from substandard measurements. This improved robustness under non-ideal conditions broadens the operational range of MP for experimentally challenging samples, while rejection of poor binding events improves the dynamic range important for detecting weak or low-affinity interactions. Additionally, the approach supports the development of automated, high-throughput pipelines by reducing reliance on expert intervention during analysis.

The application of machine learning (ML) to interference-based microscopy for nanoscopic imaging has primarily focused on low SNR particle detection to date^38,39^. These studies tested the algorithms on experimental data on the order of ∼1,000 particles. Our work shifts the emphasis toward enhancing resolving power, advancing the technique as a quantitative analytical tool, testing on over 100,000 experimental particles. A notable consequence of our approach is the interpretable feedback it provides on the distribution of event types within a given measurement. This emergent property enables users to make informed adjustments on concentration, binding conditions, or sample preparation, facilitating routine acquisition of high-quality data^18^. Such a machine vision-based filtering approach also shows great promise for application in single-molecule tracking on lipid bilayers by MP^41,42^, where extensive filtering of particles is required. MP detection and resolution remains limited by an uncharacterised, speckle-like dynamic background that emerges at high averaging^43^. Without a clear understanding of this phenomenon, our ability to simulate and surpass the limits of MP using supervised learning will remain limited.

This work represents a significant step toward integrating deep learning into quantitative mass photometry. Owing to its lightweight, thumbnail-based architecture, our network trains rapidly and lends itself to transfer learning and adaptation across local, individual instruments. We invite the community to retrain and refine the model on their own data to accelerate broader adoption. Notably, our framework integrates with established particle detection pipelines, suggesting exciting opportunities for synergy with emerging ML-based methods that jointly enhance sensitivity and resolution. While the theoretical resolution limit of MP remains at ∼5 kDa FWHM under current optical constraints^43^, our approach offers a practical path toward closing this gap. In combination with advances in detection, particle fitting, analytical corrections, and next-generation microscope designs^44,45^, such developments promise to unlock the full potential of mass photometry as a robust, high-precision tool for advanced biomolecular quantification in solution.

## Methods

### Sample Preparation

Bovine serum albumin (BSA; lyophilised powder, Sigma-Aldrich), Anti-PSMC6 antibody [p42-23] (Abcam) and protein A from *Staphylococcus aureus* (lyophilised powder, Sigma-Aldrich) were diluted to 10 – 60 µM stock solutions in Dulbecco’s PBS. Dynamin-ΔPRD samples were prepared as detailed in Foley and Kushwah et. al^42^. The expression and purification of HSP27 (HspB1, bird) were performed as previously described^46^.

### Mass Photometry measurement

All measurements followed the procedures outlined in the established protocol^18^. Briefly, we performed all measurements using a Two^MP^ instrument (Refeyn) at room temperature. We cleaned No. 1.5 coverslips (50 x 24 mm, VWR) sequentially by 5-minute sonication in milli-Q water, isopropanol, and milli-Q water, and dried using nitrogen. Grace Bio-Labs reusable CultureWell™ gaskets (3 mm diameter × 1 mm depth) contained the sample. We dispensed 4 µL of buffer to the gasket and focused the instrument with the autofocus function. We prepared working solutions by diluting samples to a final concentration of 1–100 nM and incubated for 2 minutes before measurement. Finally, we added a volume of 16 µL of the working solution, aspirated, and initiated measurement.

We recorded 60 second measurements with the small field of view setting (29.8 µm^2^) using the Acquire^MP^ software (Refeyn). Frame binning was set to 3, and data was acquired at 726 fps with an exposure time of 1.3 ms. Subsequent data analysis consisted of image pre-processing, particle picking, fitting, classification and mass histogram production. We performed these analyses using custom python scripts. Detailed information on particle classification is given below. Briefly, we performed particle picking by applying a t-test-based filter to the raw signal to detect step changes corresponding to particle landings. We further refined the selection using a radial symmetry-based filter on the ratiometric images. Unless stated otherwise, we conducted all analyses with an integration time of 40 ms (*n_avg_* = 10). To quantify the event contrast, we fitted an experimental point spread function (ePSF) to the detected particles; details of its generation are provided below. For mass calibration, we measured Dynamin-ΔPRD alongside each experimental set. Using the calibrated mass values and fitted events, we constructed mass histograms to represent the distribution for each sample.

### Training Data Generation

We generated training data by simulating particles landing on an experimental background. An experimental point spread function (ePSF) modelled the landing particles. We derived the ePSF model from the event thumbnails of the dimer species (180 kDa) of Dynamin-ΔPRD (3,302 events from 5 independent measurements), selected due to its sufficient SNR and abundance. We interpolated and centred thumbnails on the central peak, retaining the top 80% of thumbnails with high similarity scores to average and generate a representative ePSF model for the instrument.

*Optimal event simulation (binders):* To simulate stable, optimal binding, we randomly generated a single landing event (*x_i_, y_i_, t_i_*) on top of a clean buffer movie and added it to all subsequent frames. We generated the mass of the event (30 kDa ≤ *m_i_* < 800 kDa) to form the distribution shown in **Fig. 3A**. After ratiometrically processing the movie (*n_avg_* = 10), we extracted a thumbnail (40 frames x 17 px x 17 px) centred around the event. We then discarded the simulated movie and repeated the process with a clean, randomly selected buffer for every simulated event.

*Suboptimal event simulation:* We introduced additional protein dynamics to simulate sub-optimally binding events. By limiting the addition of the landing event for *t_u_* frames, the simulation accounted for *unbinding events*. The value of *t_u_*, drawn from a normal distribution with a mean of 10 and a standard deviation of 5, truncated between 0 and 20 frames. To simulate *rolling events*, the model added a landing event that moved in a singular direction for *t_r_* frames at a speed *v_r_*. The value of *t_r_* came from a uniform distribution ranging from 4 to 8 frames, while the model drew *v_r_* from a uniform distribution between 1.7 nm ms⁻¹ and 18.6 nm ms⁻¹. Upon completing its motion, the rolling event either remained stationary or detached from the surface, with a 0.75 probability of unbinding. The model simulated *neighbouring events* by introducing an optimal binder, followed by adding *N_n_* other binders randomly positioned within the event thumbnail. The value of *N_n_* ranged from 1 to 19 with equal probability. *Wobbling events* simulated proteins that followed a random walk, with its direction and speed (*v_w_*) updated at each frame over its wobbling duration (*t_w_*). The value of *t_w_* was drawn from a uniform distribution between 2 and 15 frames, while *v_w_* was sampled from a normal distribution with a mean of 8.5 nm ms^-1^ and a standard deviation of 3.4 nm ms^-1^. After wobbling, the protein either remained stationary or detached from the surface with a 0.5 probability. **Table S1** summarises all simulation parameters. **Fig. S1** depicts the effects of varying simulation parameters.

25,000 simulated events composed the training dataset. We equally represented all event classes for balanced representation. **Fig. 3A** displays the mass distribution of the training data, with a bias towards the low SNR regime (30 kDa ≤ *m* < 100 kDa) to compensate for the increased prediction uncertainty in this range.

### Model Architecture

**Fig. 3C** illustrates the 3D-ResNet based deep learning model^21,40^ used for classification. The model receives a 3D spatiotemporal thumbnail input, centred on the detected landing event, with dimensions of 40 frames x 17 pixels x 17 pixels. The model consists of an initial 3D convolutional layer (7 x 3 x 3) followed by batch normalisation, a ReLU activation layer and max pooling (3 x 3 x 3). This is followed by 4 residual layers consisting of 3, 4, 6 and 3 convolutional blocks, respectively. Residual connections with identity downsampling were implemented after the first convolutional block in each residual layer. Each convolutional block consisted of three 3D convolutional layers followed by batch normalisation and a final ReLU activation layer. After the final residual layer, global average pooling and two fully connected layers progressively reduce the feature dimensions to 5 in the final output layer.

### Model Training

Data pre-processing consisted of removing any thumbnails that contained NaN values. We performed standardisation on a per-thumbnail basis using a z-score

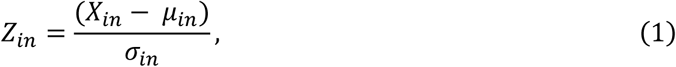

where 𝑍_𝑖𝑛_ represents the standardised thumbnail, 𝑋_𝑖𝑛_ is the input thumbnail, and 𝜇_in_ and σ_in_ are the mean and standard deviation of the input thumbnail. This standardisation generalised mass bias during training and improved model performance across the entire mass range. We split the dataset 80:20 for training and validation, with a batch size of 10 and random shuffling.

Model parameters were randomly initialised prior to training. We trained and optimised the model using a cross-entropy loss function and stochastic gradient descent at a learning rate of 0.1. A learning rate scheduler reduced the learning rate by a factor of 0.1 if the validation loss plateaued for 5 consecutive epochs. Gradients were clipped to a maximum norm value of 0.5 to prevent exploding gradients in deeper layers. Training occurred for 20 epochs. The validation accuracy plateaued at 94.5% after 15 epochs, at which point the model was accepted to avoid overfitting to the training data. The total training time was 64 minutes. Given the size of the task, we performed training on an NVIDIA GeForce MX450 at an acceptable computational cost.

The model was trained to evaluate thumbnails ratiometrically processed with an integration time of 40 ms (*n_avg_* = 10). A second model was also trained, designed to perform on data processed with an integration time of 160 ms (*n_avg_* = 40, corresponding to 80 frames x 17 px x 17 px thumbnails).

### Evaluation and Validation

#### Confusion matrix

A confusion matrix was calculated from the validation dataset and shown in **Fig. 3F**.

#### Receiver Operating Characteristic (ROC) curves

**Fig. 3H** shows the ROC curves separated by event mass and class, with **Table S2** reporting the AUC. We performed this analysis for the entire validation set of 5,000 events, covering a mass range from 30 kDa to 800 kDa. To further assess performance across three specific mass ranges (30 kDa to 55 kDa, 55 kDa to 100 kDa, and 100 kDa to 800 kDa), we simulated and evaluated 1,500 additional events within each range, distributing them evenly across all event classes. We used the model as a binary classifier to assess the performance across each class. The output of the trained model was passed through a sigmoid function to obtain classification scores, and ROC curves were generated by varying the decision threshold for the target class.

#### Experimental data evaluation

We extracted and standardised 3D thumbnails (40 frames × 17 pixel × 17 pixel) centred on each detected event. The deep learning model assigned class scores to each thumbnail. We determined an optimal classification threshold for the binder class and retained only events with scores above this threshold. To assess the improvement in resolution performance, we determined the mass σ by fitting a Gaussian curve to clearly resolved mass peaks. For partially resolved peaks, such as those observed with HSP27 – bird, we calculated the valley-to-peak ratio (VPR) for each peak set by taking the ratio between the minimum point in the valley and the height of the smaller peak.

#### Varying event density

We evaluated the effect of event filtering on mass σ for increasingly dense movies using simulated dynamin-ΔPRD landing assays with optimal binding. We quantified the results based on the tetramer species (360 kDa). We simulated each density with 5 repeats. To specifically isolate the effects of event density, we simulated each movie with a constant oligomeric distribution. We applied a binder score threshold of 0.8 to retain the most optimal events, recognising that most events had neighbouring signals.

#### Varying integration time

We evaluated the effect of event filtering on mass σ for various integration times using dynamin-ΔPRD. We quantified the results based on the tetramer species (360 kDa). To perform classification, we trained two models at distinct integration times: 40 ms (*n_avg_* = 10) and 160 ms (*n_avg_* = 40). We then varied the integration time of the detected and classified events for subsequent analysis. For integration times below 160 ms, we used classifications from the first model; for times above 160 ms, we used classifications from the second model. This was repeated for 5 independent measurements.

#### Low mass proteins

To evaluate performance on low mass proteins, we tested the model on protein A. To prevent cut-off by the detection algorithm, we set permissive filter thresholds: Filter 1 = 0.01, Filter 2 = 0.15. We performed classification with a high binder score threshold of 0.997 to reduce the false positive rate in the low mass regime. After classification, we applied a nearest neighbour filter to remove multiple detections of the same landing event. This analysis was carried out across a titration series, with a minimum of four repeats per concentration.

#### Validation

We validated the mass distribution of the antibody measurements against SDS-PAGE analysis and the mass distribution of the HSP27 measurements using native mass spectrometry^46^. We benchmark the performance on the HSP27 measurements against analysis from the Discover^MP^ software across various Filter 1 and Filter 2 settings, with results summarised in **Table S3** and **Fig. S5**. Additionally, classification outcomes were further evaluated by comparison to the fit residuals (𝑟), which were calculated as the square root of the sum of squared residuals between the measured events X(i, j) and fitted events X_fit_(i, j), as illustrated in **Fig. S4**,

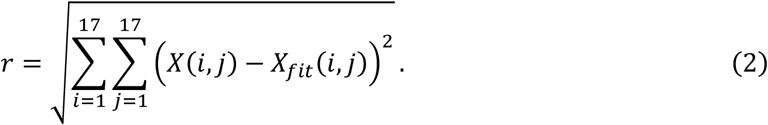

## Funding Sources

The work was supported by European Research Council (ERC) Consolidator Grant PHOTOMASS 819593 (P.K., K.I., J.S.P.), Engineering and Physical Sciences Research Council (EPSRC) Leadership Fellowship EP/T03419X/1 (P.K., J.C.T.,S.Tu.), Schmidt Sciences, LLC (J.C.T.), Clarendon Scholarship (D.S.), EPRSC Doctoral Training Partnership (J.B.) and Biotechnology and Biological Sciences Research Council BB/W00349X/1 (J.L.P.B., P.K., S.Th.), Leverhulme Trust RPG-2021-246 (D.S.).

## Supporting information

Supplementary Figures accompanying the manuscript.

## Tables

**Table S1.**
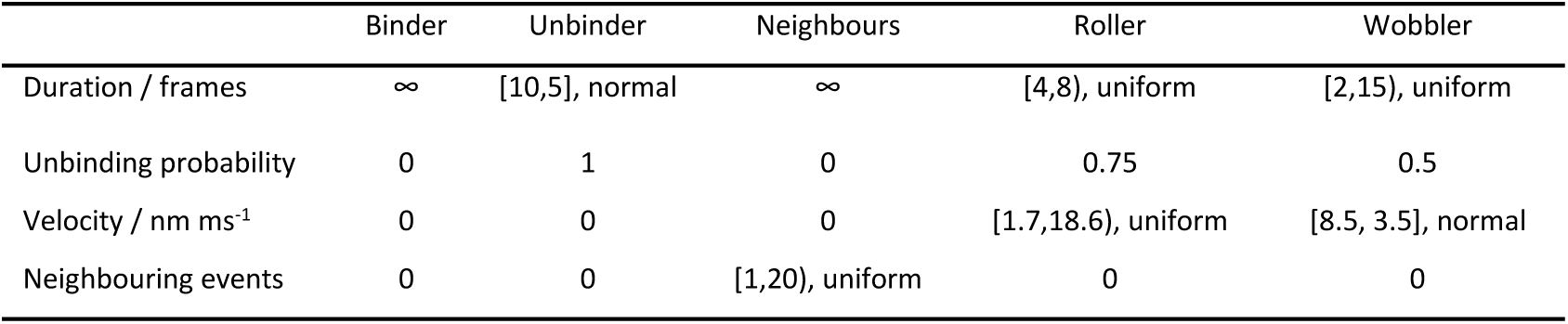
Simulation parameters for training data by event class. Event masses were randomly generated between 30 and 800 kDa, with a bias towards the low mass range below 100 kDa.

**Table S2.** Area under the curve (AUC) for validation dataset (30–800 kDa, 5,000 events) and three additional datasets subdivided by mass range (1,500 events each), with diagnostic performance separated by class.

|  | Binder | Unbinder | Neighbours | Roller | Wobbler |
| --- | --- | --- | --- | --- | --- |
| Validation | 0.985 | 0.999 | 0.995 | 0.996 | 0.985 |
| 30 ≤ m (kDa) < 55 | 0.841 | 0.995 | 0.882 | 0.968 | 0.868 |
| 55 ≤ m (kDa) < 100 | 0.932 | 0.999 | 0.978 | 0.998 | 0.950 |
| 100 ≤ m (kDa) < 800 | 0.998 | 0.999 | 0.995 | 1.00 | 0.995 |

**Table S3.** Resolution of HSP27 bird sample, quantified by the valley-to-peak ratio of partially resolved peak pairs. Resolution was assessed using standard Discover^MP^ analysis settings (*n_avg_ = 5, Threshold 1 = 1.20, Threshold 2 = 0.25*), user-optimised Discover^MP^ settings (*n_avg_ = 10, Threshold 1 = 2.60, Threshold 2 = 0.25*), and custom Python analysis (n_avg_ = 10) before and after optimal event filtering. *m* indicates the number of independent repeat measurements, and *n* represents the estimated total number of events for each partially resolved peak pair. The mass of the HSP27 monomer is approximately 22.8 kDa.

| Sample | Valley-to-peak ratio |  |  |  | Resolving Power<br>(30% valley) |
| --- | --- | --- | --- | --- | --- |
|  | DMP<br>Standard | DMP<br>Optimised | Before event<br>filtering | After event<br>filtering |  |
| 6-mer, 8-mer<br>(m = 4, n = 1271) | 0.6 ± 0.1 | 0.5 ± 0.1 | 0.69 ± 0.05 | 0.06 ± 0.05 | 3.5 |
| 8-mer, 10-mer<br>(m = 4, n = 1592) | 0.38 ± 0.01 | 0.27 ± 0.05 | 0.42 ± 0.05 | 0.09 ± 0.01 | 4.5 |
| 10-mer, 12-mer<br>(m = 4, n = 2057) | 0.51 ± 0.03 | 0.30 ± 0.04 | 0.38 ± 0.03 | 0.25 ± 0.03 | 5.5 |
| 12-mer, 14-mer<br>(m = 4, n = 2062) | 0.41 ± 0.02 | 0.33 ± 0.03 | 0.38 ± 0.02 | 0.23 ± 0.04 | 6.5 |
| 14-mer, 16-mer<br>(m = 4, n = 1710) | 0.46 ± 0.03 | 0.43 ± 0.05 | 0.5 ± 0.1 | 0.3 ± 0.1 | 7.5 |

## Notes

### Competing Interest Statement

The authors declare the following competing interests: P.K. is a nonexecutive director, shareholder of and consultant to Refeyn Ltd., J.L.P.B. is a shareholder of and consultants to Refeyn Ltd.

## References

1 Young, G. et al. Quantitative mass imaging of single biological macromolecules. Science 360, 423-+ (2018). 10.1126/science.aar5839

2 Sonn-Segev, A. et al. Quantifying the heterogeneity of macromolecular machines by mass photometry. Nat Commun 11 (2020). https://doi.org/ARTN 1772 10.1038/s41467-020-15642-w

3 Gizardin-Fredon, H. et al. Denaturing mass photometry for rapid optimization of chemical protein-protein cross-linking reactions. Nat Commun 15, 3516 (2024).

4 Soltermann, F. et al. Quantifying Protein-Protein Interactions by Molecular Counting with Mass Photometry. Angew Chem Int Edit 59, 10774–10779 (2020). 10.1002/anie.202001578

5 Wu, D. & Piszczek, G. Measuring the affinity of protein-protein interactions on a single-molecule level by mass photometry. Anal Biochem 592 (2020). https://doi.org/ARTN 113575 10.1016/j.ab.2020.113575

6 Haüssermann, K., Young, G., Kukura, P. & Dietz, H. Dissecting FOXP2 Oligomerization and DNA Binding. Angew Chem Int Edit 58, 7662–7667 (2019). 10.1002/anie.201901734

7 Kissling, V. M. et al. Mre11-Rad50 oligomerization promotes DNA double-strand break repair. Nat Commun 13 (2022). https://doi.org/ARTN 2374 10.1038/s41467-022-29841-0

8 Zinder, J. C. et al. Shelterin is a dimeric complex with extensive structural heterogeneity. P Natl Acad Sci USA 119 (2022). https://doi.org/ARTN e2201662119 10.1073/pnas.2201662119

9 Young, J. W. et al. Characterization of membrane protein interactions by peptidisc-mediated mass photometry. Iscience 27 (2024). https://doi.org/ARTN 108785 10.1016/j.isci.2024.108785

10 Sendker, F. L. et al. Emergence of fractal geometries in the evolution of a metabolic enzyme. Nature (2024). 10.1038/s41586-024-07287-2

11 Cole, D., Young, G., Weigel, A., Sebesta, A. & Kukura, P. Label-Free Single-Molecule Imaging with Numerical-Aperture Shaped Interferometric Scattering Microscopy. Acs Photonics 4, 211–216 (2017). 10.1021/acsphotonics.6b00912

12 Piliarik, M. & Sandoghdar, V. Direct optical sensing of single unlabelled proteins and super-resolution imaging of their binding sites. Nat Commun 5 (2014). https://doi.org/ARTN 4495 10.1038/ncomms5495

13 Murray, K. K. Resolution and Resolving Power in Mass Spectrometry. J Am Soc Mass Spectr 33, 2342–2347 (2022). 10.1021/jasms.2c00216

14 Infante, H. G. et al. Glossary of methods and terms used in analytical spectroscopy (IUPAC Recommendations 2019). Pure Appl Chem 93, 647–776 (2021). 10.1515/pac-2019-0203

15 D. N. Brett, G. W. E., and D. A. Skoog. Compendium of Analytical Nomenclature: Definitive Rules 1997. 3rd Edition edn, (Blackwell Science for IUPAC, 1998).

16 Kühlbrandt, W. The Resolution Revolution. Science 343, 1443–1444 (2014). 10.1126/science.1251652

17 Sülzle, J., Elfeky, L. & Manley, S. Surface passivation and functionalisation for mass photometry. J Microsc-Oxford 295, 14–20 (2024). 10.1111/jmi.13302

18 Kratochvíl, J. et al. Best practice mass photometry: A guide to optimal single molecule mass measurement. bioRxiv, 2024.2012.2003.624087 (2024). 10.1101/2024.12.03.624087

19 LeCun, Y., Bengio, Y. & Hinton, G. Deep learning. Nature 521, 436–444 (2015). 10.1038/nature14539

20 Krizhevsky, A., Sutskever, I. & Hinton, G. E. ImageNet Classification with Deep Convolutional Neural Networks. Commun Acm 60, 84–90 (2017). 10.1145/3065386

21 He, K. M., Zhang, X. Y., Ren, S. Q. & Sun, J. Deep Residual Learning for Image Recognition. Proc Cvpr Ieee, 770–778 (2016). 10.1109/Cvpr.2016.90

22 Lecun, Y., Bottou, L., Bengio, Y. & Haffner, P. Gradient-based learning applied to document recognition. P Ieee 86, 2278–2324 (1998). Doi 10.1109/5.726791

23 Vaswani, A., et al. Attention Is All You Need. Adv Neur In 30 (2017).

24 Midtvedt, B. et al. Quantitative digital microscopy with deep learning. Appl Phys Rev 8 (2021). https://doi.org/ARTN 011310 10.1063/5.0034891

25 Pineda, J. et al. Geometric deep learning reveals the spatiotemporal features of microscopic motion. Nat Mach Intell 5, 71-+ (2023). 10.1038/s42256-022-00595-0

26 Belthangady, C. & Royer, L. A. Applications, promises, and pitfalls of deep learning for fluorescence image reconstruction. Nat Methods 16, 1215–1225 (2019). 10.1038/s41592-019-0458-z

27 Zhou, S. K. et al. A Review of Deep Learning in Medical Imaging: Imaging Traits, Technology Trends, Case Studies With Progress Highlights, and Future Promises. P Ieee 109, 820–838 (2021). 10.1109/Jproc.2021.3054390

28 Huertas-Company, M. & Lanusse, F. The Dawes Review 10: The impact of deep learning for the analysis of galaxy surveys. Publ Astron Soc Aust 40 (2023). https://doi.org/ARTN e01 PII S1323358022000558 10.1017/pasa.2022.55

29 Weigert, M. et al. Content-aware image restoration: pushing the limits of fluorescence microscopy. Nat Methods 15, 1090-+ (2018). 10.1038/s41592-018-0216-7

30 Möckl, L., Roy, A. R., Petrov, P. N. & Moerner, W. E. Accurate and rapid background estimation in single-molecule localization microscopy using the deep neural network BGnet. P Natl Acad Sci USA 117, 60–67 (2020). 10.1073/pnas.1916219117

31 Yang, T., Luo, Y., Ji, W. & Yang, G. Advancing biological super-resolution microscopy through deep learning: a brief review. Biophys Rep 7, 253–266 (2021). 10.52601/bpr.2021.210019

32 Gomariz, A. et al. Modality attention and sampling enables deep learning with heterogeneous marker combinations in fluorescence microscopy. Nat Mach Intell 3, 799-+ (2021). 10.1038/s42256-021-00379-y

33 Young, A., Röst, H. & Wang, B. Tandem mass spectrum prediction for small molecules using graph transformers. Nat Mach Intell 6 (2024). 10.1038/s42256-024-00816-8

34 Yu, Y. H. & Li, M. Towards highly sensitive deep learning-based end-to-end database search for tandem mass spectrometry. Nat Mach Intell 7 (2025). 10.1038/s42256-024-00960-1

35 Giri, N., Roy, R. S. & Cheng, J. L. Deep learning for reconstructing protein structures from cryo-EM density maps: Recent advances and future directions. Curr Opin Struc Biol 79 (2023). https://doi.org/ARTN 102536 10.1016/j.sbi.2023.102536

36 Huang, Y., Zhu, C. G., Yang, X. K. & Liu, M. H. High-resolution real-space reconstruction of cryo-EM structures using a neural field network. Nat Mach Intell 6 (2024). 10.1038/s42256-024-00870-2

37 Matsumoto, S. et al. Extraction of protein dynamics information from cryo-EM maps using deep learning. Nat Mach Intell 3 (2021). 10.1038/s42256-020-00290-y

38 Boyle, M. J., Goldman, Y. E. & Composto, R. J. Enhancing Nanoparticle Detection in Interferometric Scattering (iSCAT) Microscopy Using a Mask R-CNN. J Phys Chem B 127, 3737–3745 (2023). 10.1021/acs.jpcb.3c00097

39 Dahmardeh, M., Dastjerdi, H. M., Mazal, H., Köstler, H. & Sandoghdar, V. Self-supervised machine learning pushes the sensitivity limit in label-free detection of single proteins below 10 kDa. Nat Methods 20, 442-+ (2023). 10.1038/s41592-023-01778-2

40 Hara, K., Kataoka, H. & Satoh, Y. Learning Spatio-Temporal Features with 3D Residual Networks for Action Recognition. Ieee Int Conf Comp V, 3154-3160 (2017). 10.1109/Iccvw.2017.373

41 Heermann, T., Steiert, F., Ramm, B., Hundt, N. & Schwille, P. Mass-sensitive particle tracking to elucidate the membrane-associated MinDE reaction cycle. Nat Methods 18, 1239-+ (2021). 10.1038/s41592-021-01260-x

42 Foley, E. D. B., Kushwah, M. S., Young, G. & Kukura, P. Mass photometry enables label-free tracking and mass measurement of single proteins on lipid bilayers. Nat Methods 18, 1247-+ (2021). 10.1038/s41592-021-01261-w

43 Becker, J. et al. A Quantitative Description for Optical Mass Measurement of Single Biomolecules. Acs Photonics 10, 2699–2710 (2023). 10.1021/acsphotonics.3c00422

44 Iqbal, K., Thiele, J. C., Pfitzner, E. & Kukura, P. Enhanced Interferometric Imaging by Rotating Coherent Scattering Microscopy. Acs Photonics (2025). 10.1021/acsphotonics.5c00123

45 Asor, R., Loewenthal, D., van Wee, R., Benesch, J. L. & Kukura, P. Mass Photometry. Annu Rev Biophys 54, 379–399 (2025).

46. aman, D. Quantitative Description of Co-Assembly and Evolution of Small Heat-Shock Proteins DPhil thesis thesis, University of Oxford, (2021).

