## Supplementary Figures accompanying the manuscript. for "Deep learning-based event classification of mass photometry data for optimal mass measurement at the single-molecule level"


### Supporting Information

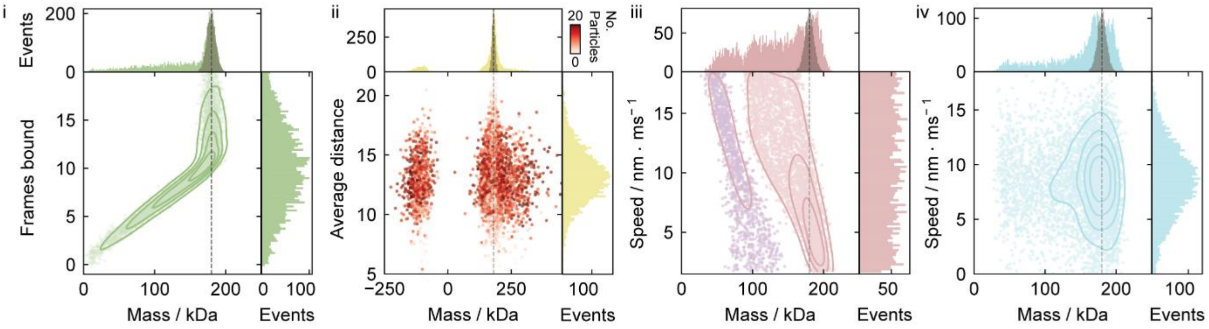


**Fig S1** – The mass broadening effect of varying the simulated dynamics of suboptimal 180 kDa events. (i) Effect of transient unbinding as a function of frames bound. (ii) Effects of particle density and proximity in high event density thumbnails. (iii) Effect of rolling velocity. (iv) Effect of wobbling velocity.

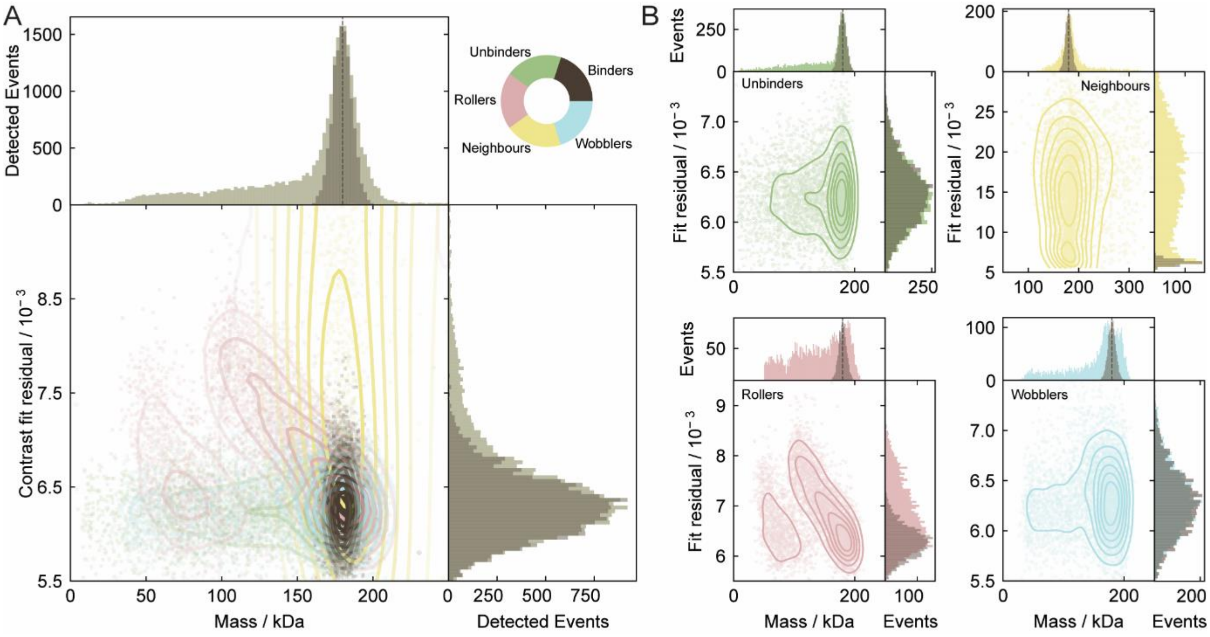


**Fig S2** – Investigating the effect of suboptimal binding on mass photometry through simulation. (A) Contour plot depicting 25,000 simulated 180 kDa events, evenly distributed across each event class with different simulation parameters. Mass and residual histograms for all combined classes are shown (light), with the normalised histograms for the binder class overlaid (dark). Mass broadening is observed with a shoulder and increased base line noise extending across the low mass range. (B) Mass-residual contour plot separated by event class, demonstrating the mass broadening effect of transient behaviours such as rapid unbinding or wobbling, which cannot be distinguished using the 2D information in fit residuals alone.

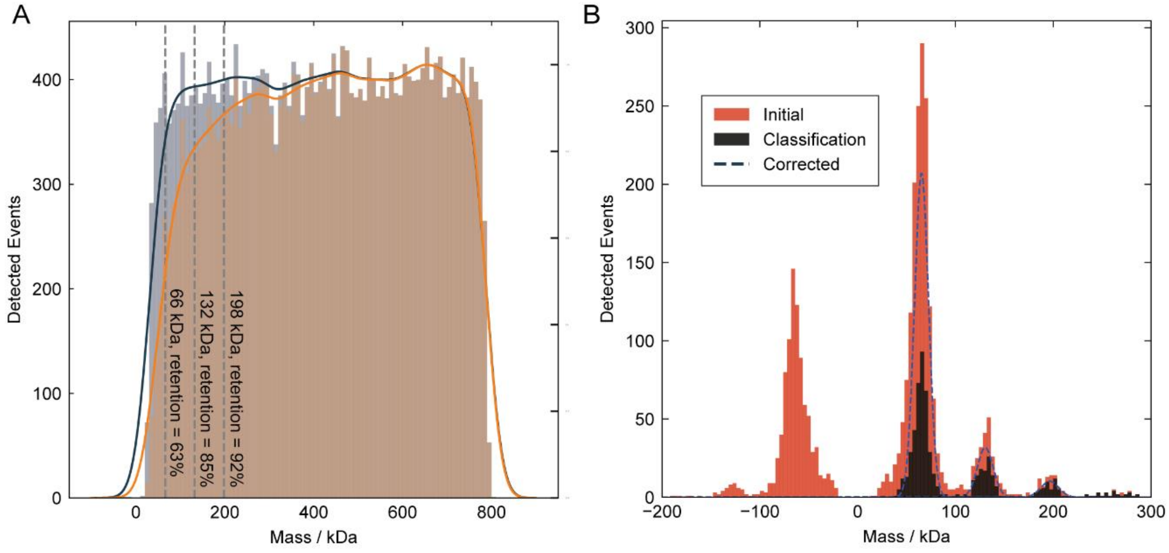


**Fig. S3 –** (A) Misclassification rates of optimal binding events across varying masses. Histogram of 29,901 simulated optimal events (slate blue) and retained events post-classification (orange). A normalised KDE plot overlays the histograms, used to calculate the retention rate of optimal events by mass. (B) Corrected BSA histogram, accounting for misclassification of optimal events.

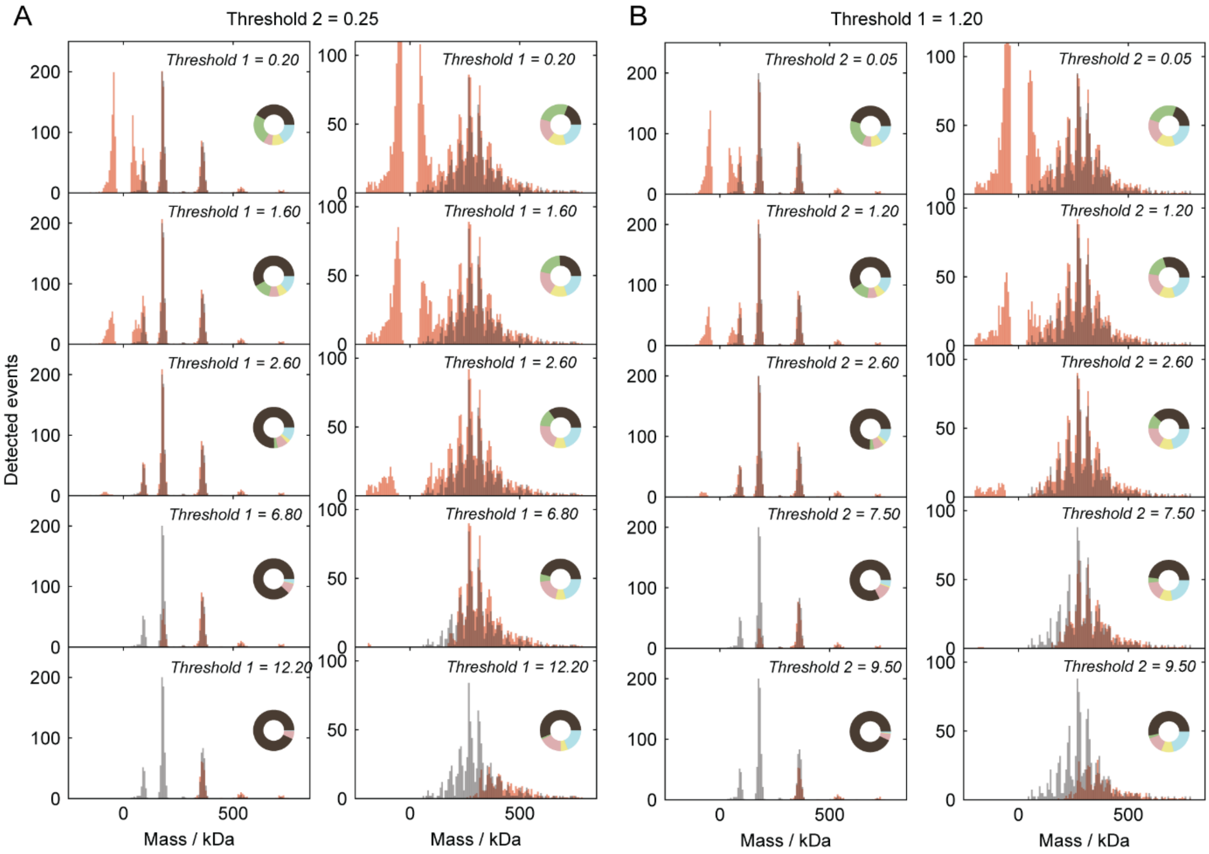


**Fig S4** – The effect of adjusting Filter 1 and Filter 2 settings in Discover$^{MP}$. Demonstrated effects for Dynamin-ΔPRD and HSP27 bird samples, illustrating how arbitrary changes can alter the observed mass distribution. The Discover$^{MP}$ analysis is shown in red, with the classification filtering analysis scaled and overlaid in grey. Pie chart insets display the distribution of event classes in the Discover$^{MP}$ analyses. (A) Effect of varying Filter 1 with Filter 2 fixed at 0.25. (B) Effect of varying Filter 2 with Filter 1 fixed at 1.20.

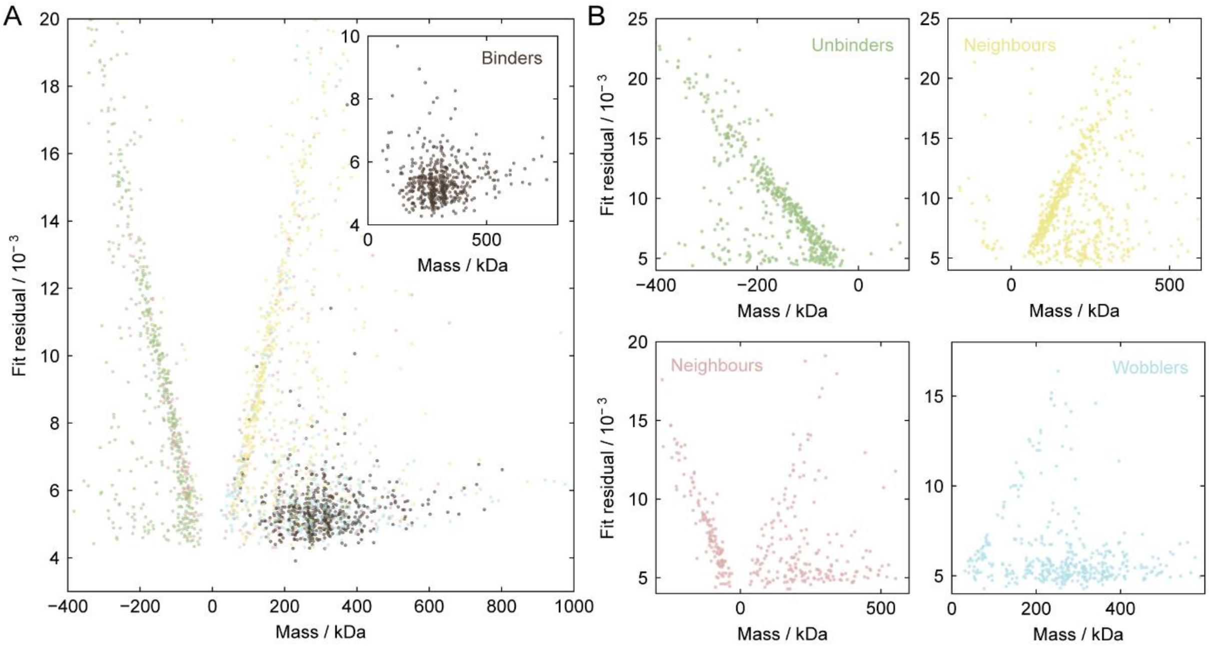


**Fig S5**– Contrast fit residuals for HSP27. (A) Combined fit residuals for all landing events, with an inset displaying the fit residuals for the optimal binder class. (B) Fit residuals separated for the suboptimal event classes.

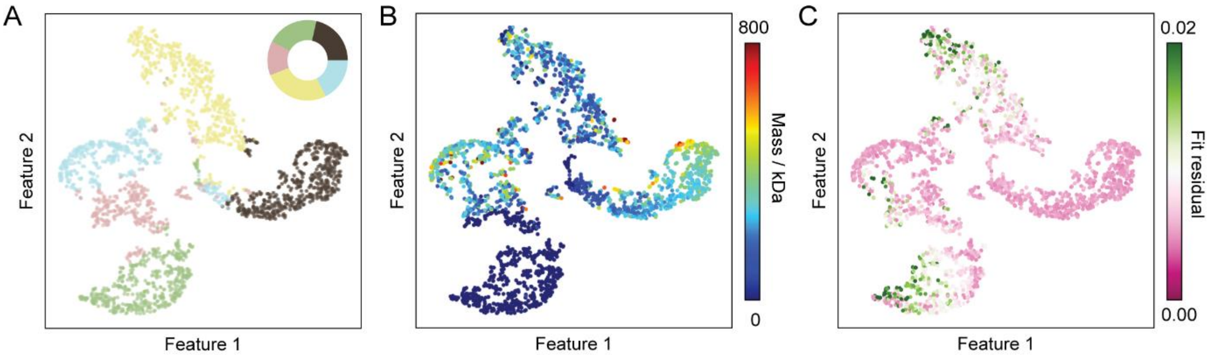


**Fig S6** – t-SNE feature space representation for HSP27. (A) Event class distribution. (B) Mass distribution. (C) Fit residual distribution.

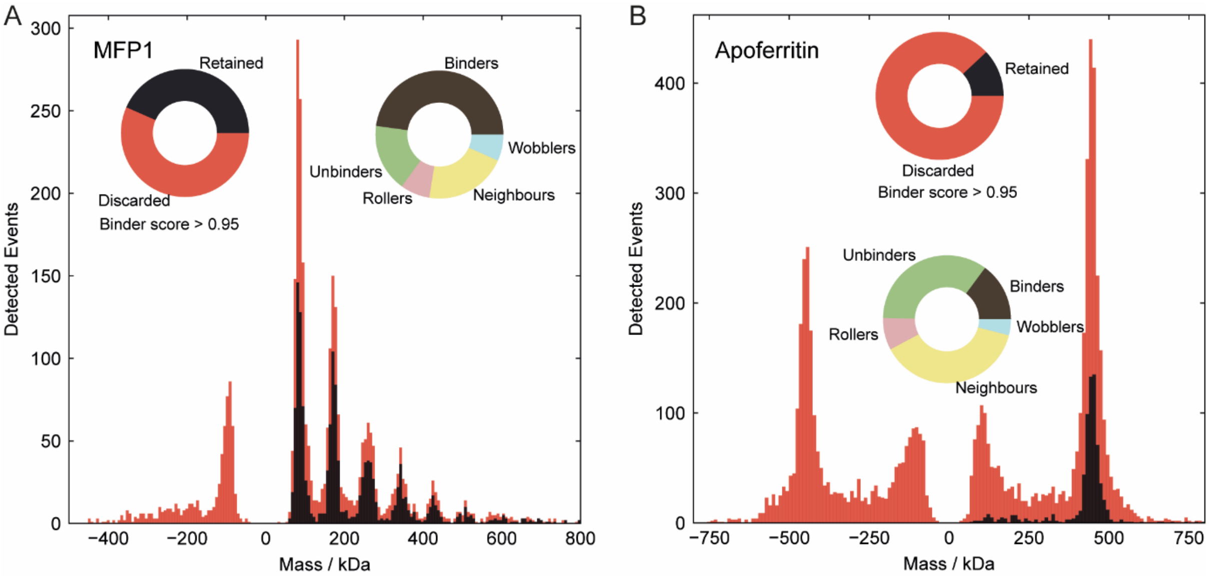


**Fig S7** – Identifying optimal single-molecule landing events for massference-p1 and apoferritin.

**Supplementary Video 1 –** Thumbnails of landing events from the HSP27 dataset, categorised into optimal and suboptimal events. Each thumbnail is individually z-scaled (colour bar) to optimise image contrast.
